# Telomere-to-telomere African wild rice (*Oryza longistaminata*) reference genome reveals segmental and structural variation

**DOI:** 10.1101/2024.09.05.611405

**Authors:** Xuanmin Guang, Jingnan Yang, Shilai Zhang, Fei Guo, Linzhou Li, Xiaoping Lian, Tao Zeng, Chongyang Cai, Fushu Liu, Zhihao Li, Yangzi Hu, Dongming Fang, Weiming He, Sunil Kumar Sahu, Wangsheng Li, Haorong Lu, Yuxiang Li, Huan Liu, Xun Xu, Ying Gu, Fengyi Hu, Yuliang Dong, Tong Wei

## Abstract

Rice (*Oryza sativa*) is one of the most important staple food crops worldwide, and its wild relatives serve as an important gene pool in its breeding. Compared with cultivated rice species, African wild rice (*Oryza longistaminata*) has several advantageous traits, such as resistance to increased biomass production, clonal propagation via rhizomes, and biotic stresses. However, previous *O. longistaminata* genome assemblies have been hampered by gaps and incompleteness, restricting detailed investigations into their genomes. To streamline breeding endeavors and facilitate functional genomics studies, we generated a 343-Mb telomere-to-telomere (T2T) genome assembly for this species, covering all telomeres and centromeres across the 12 chromosomes. This newly assembled genome has markedly improved over previous versions. Comparative analysis revealed a high degree of synteny with previously published genomes. A large number of structural variations were identified between the *O. longistaminata* and *O. sativa*. A total of 2,466 segmentally duplicated genes were identified and enriched in cellular amino acid metabolic processes. We detected a slight expansion of some subfamilies of resistance genes and transcription factors. This newly assembled T2T genome of *O. longistaminata* provides a valuable resource for the exploration and exploitation of beneficial alleles present in wild relative species of cultivated rice.

## Introduction

Rice stands as one of the world’s most essential crops, serving as a staple food source for over half of the global population [1]. Rice breeding is therefore critical for global food security, for which germplasms, especially wild relatives, serve as an important gene pool. *Oryza longistaminata* (2x=2n=12), an AA genome type, thrives predominantly in the tropical regions of Western Africa, often in proximity to freshwater sources and swampy areas[2]. Although it is rarely used for human consumption, this species possesses a variety of beneficial traits. Notably, it is resistant to bacterial blight, which is linked to the *Xa21* locus[3]. Furthermore, *O. longistaminata* exhibits perennial growth and an exceptional capacity for biomass production and based on which much efforts have been made to transfer these beneficial alleles into those commercial rice varieties. In addition to contributions to breeding endeavors, *O. longistaminata* serves as a vital subject of study for investigating the genetic foundations and developmental aspects of rhizomes[4].

The assembly of a complete plant genome provides a solid basis for functional genomics investigations and facilitates the identification of candidate genes using traditional mapping techniques. Despite the publication of several genome assembly versions, limitations stemming from sequencing technology and the intricate organization of the genome have left certain complex regions underrepresented in this reference[5, 6]. To achieve a more comprehensive representation of this fundamental reference genome, we employed a hybrid assembly strategy using Pacbio HiFi and CycloneSEQ ultra-long reads (a new single-molecule sequencer from MGI) [7] to generate backbone contigs. These contigs were subsequently scaffolded into a chromosome-level assembly with the assistance of Hi-C datasets. In addition, gap filling was executed to resolve any remaining gaps. To this end, we generated a telomere-to-telomere (T2T) assembly for *O. longistaminata*, which could serve as a valuable genomic resource for future rice research and breeding.

## Results and discussion

### Genome assembly

We initially sequenced the genome of *O. longistaminata*, generating 27.3 Gb (~ 78 × coverage) of PacBio HiFi reads, 32 Gb (~ 100 × coverage) of Hi-C paired reads, 25.6 Gb (~71.4 × coverage) of Ultra-long CycloneSEQ reads, and 21.0 Gb (~ 60 × coverage) of MGI-Seq pair end reads (Table S1). Using the *K-mer* method, we estimated the genome size of this plant to be 357 Mb,, and its heterozygosity is 1.27% (Figure S1), which is similar to the size of the previous results[6]. Using the combined data, we first assembled a genome with a size of 343 Mb and a contig N50 of 26.02 Mb. Based on the Hi-C data, we anchored 13 contigs into 12 pseudochromosomes (Table 1, Figure S2). After that, TGS-gapcloser was employed to close the remaining gaps [8]. Finally using the seven-base telomeric repeat (CCCTAAA at the 5’ end or TTTAGGG at the 3’ end) as a sequence query, we identified all the 24 telomeres for the genome (Figure S3)[9]. We then used quartets to identify the centromeric regions ranging from 0.3 to 1.8 Mb on each chromosome, and assessed the regions using Hi-C data[10]. Different methods were used to evaluate the accuracy and completeness of the assembly. First, pair-end library reads were mapped to the genome and more than 97.27% of them were aligned. Second, the BUSCO analysis indicated that the genome’s completeness reached 98.6% for the genome[11] (Table S2). Third, the LTR assembly index (LAI) value for the genome was 20.71, meeting the gold standard for genome assemblies [12]. Fourth, the calculated QV (assembly consensus quality value) using Merqury was 52.08 which indicates that the base call accuracy of the genome was higher than 99.999%[13]. As there is a previously published *O. longistaminata* genome assembly[5], we conducted a comparative gene synteny analysis on all coding sequence (CDS) levels. Subsequently, we identified a total of 28,627 syntenic CDS pairs through the genome-wide alignment. Consistent with expectations, these two genomes displayed rather high synteny, as evidenced by the pronounced central diagonal in the alignment (Figure S4). This result demonstrates the high concordance between our assembled T2T genome and the previous *O. longistaminata* assembly.

**Table 1.** Summary statistics of *O. longistaminata* genome assembly.

| Genome Feature | Value |
| --- | --- |
| Total size of assembled contigs (bp) | 343,752,306 |
| GC content | 43.02% |
| Contig N50 | 26,021,309 |
| Number of Contigs | 197 |
| Total size of assembled genomes (bp) | 331,045,917 |
| Scaffold N50 | 26,021,309 |
| Complete BUSCOs | 98.6% |
| LAI value | 20.71 |
| Repeat region | 40.73% |
| Number of gap-free chromosomes | 12 |
| Number of candidate telomeres | 24 |
| Number of candidate centromeres | 12 |
| Number of chromosomes | 12 |
| Number of protein-coding genes | 33,177 |
| Average gene length (bp) | 2439 |
| Average exon length (bp) | 261 |
| Average intron length (bp) | 384 |

### Genome annotation

Through the utilization of *de novo* and homology-based methods, we successfully identified a total of 134 Mb of repetitive sequences in the plant’s genome. These repetitive sequences make up approximately 40.73% of the entire genome. Furthermore, we observed that the repeat contents were highly consistent across the entire genome as well as among the 12 pseudochromosome sequences. (Figure 1,Table S3). In this genome, LTRs and DNA transposons were the major types of repeats, accounting for approximately 20.9% and 18.5% of the whole genome, respectively. The overall repeat content was at a moderate level, similar to the repeat content observed in other assemblies with the *Oryza* genus genomes[14].

**Figure 1.**
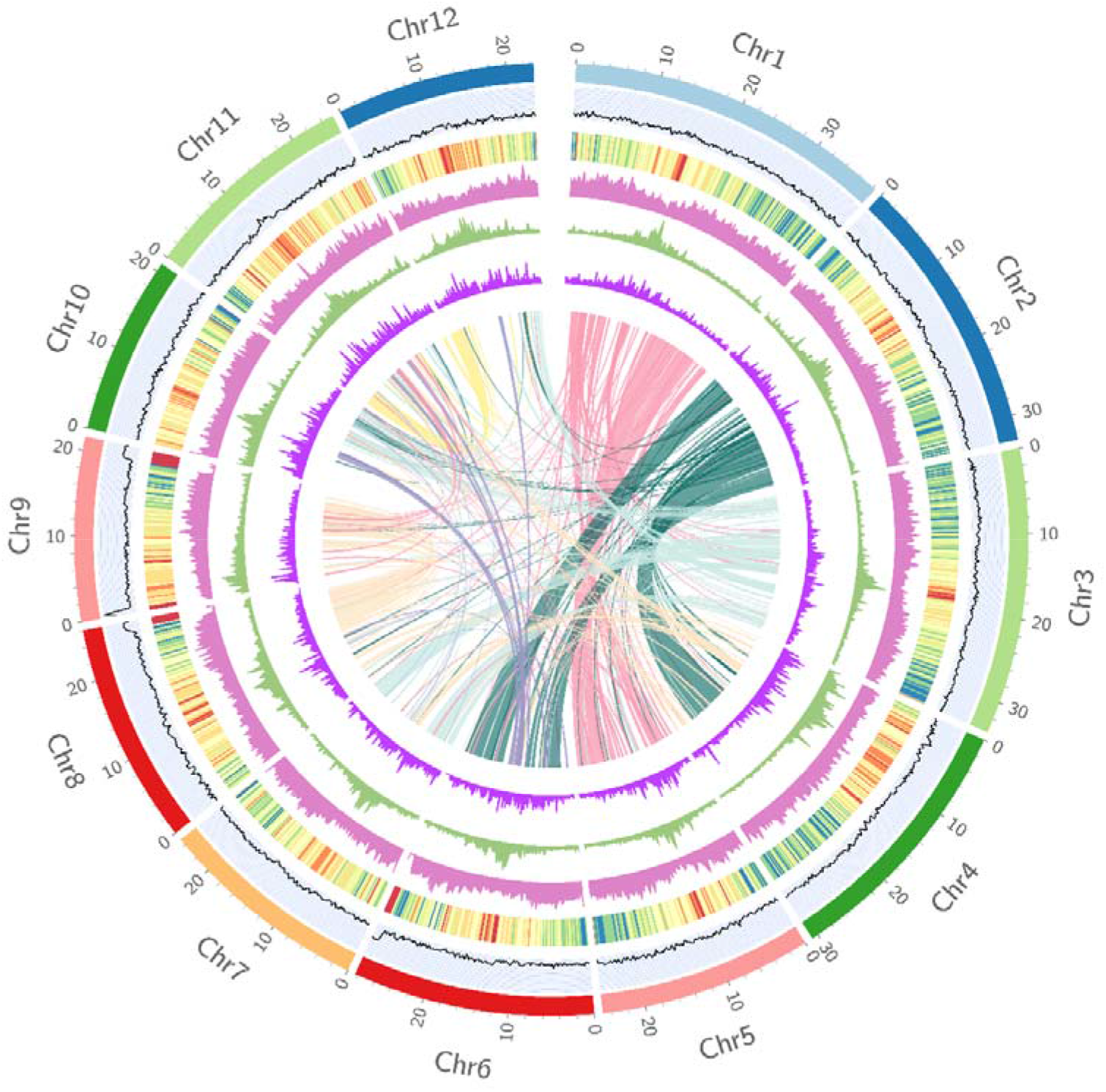
The telomere-to-telomere genome assembly of *O. longistaminata*. Genomic features of the *O. longistaminata* genome, from outer to inner, including GC percentage, protein-coding genes, repeat sequences, LTR-*Gypsy*, LTR-*Copia*. The collinear blocks are shown in the center.

The centromere region of the genome poses a significant challenge for assembly due to its high degree of repetitive sequence content[15]. To date, the centromeric sequence of the *O. longistaminata* genome has not been fully characterized, and our new T2T genome allows deeper exploration of the repeats in these regions. The results revealed a phenomenon that the centromeric regions with high densities of transposable elements and relatively low gene densities. Among the repeats of the centromeric regions, Gypsy elements were the most dominant type of LTR (Figure S5).

A total of 33,177 coding genes were predicted in this genome, with an average gene length of 2,439 bp and an average coding sequence (CDS) length of 1,138 bp (Figure 1, Table 1, Table S4). The functional analysis revealed that 95.74% of the coding genes could be annotated through publicly protein datasets (Table S5), suggesting the accuracy of gene prediction.

### Genome structural variations

We further performed genome-wide detection of putative structural variations (SVs) in the *O. sativa* genome (Figure 2, Table S6). A total of 3,738,150 single nucleotide polymorphisms (SNPs) were identified by comparing the two genomes. Among the SVs identified in our study, 204 were inversions. These inversions varied in size, ranging from 1,040 bp to 4,529,518 bp, with a median size of 6.28 kb. Additionally, we identified 11,706 duplications, and its length spanning from 999 to 134,657 bp, having a median size of 2,202 bp. Furthermore, we found 11,175 inverted duplications, spanning from 999 to 37,420 bp, feature a median size of 2162 bp. Additionally, we observed 3,077 translocations, spanning from 1,000 to 461,145 bp, with a median size of 2,093 bp. Finally, we detected 3,015 inverted translocations, covering from 999 to 70,158 bp, with a median size of 2,047 bp. These large SVs span more than 105 Mb throughout the entire genome, which indicates remarkable divergence between these two species. Gene Ontology (GO) enrichment analysis of these SV-related genes revealed that they were associated with catalytic activity, purine ribonucleotide binding, adenyl ribonucleotide binding, and telomere maintenance (Table S7).

**Figure 2.**
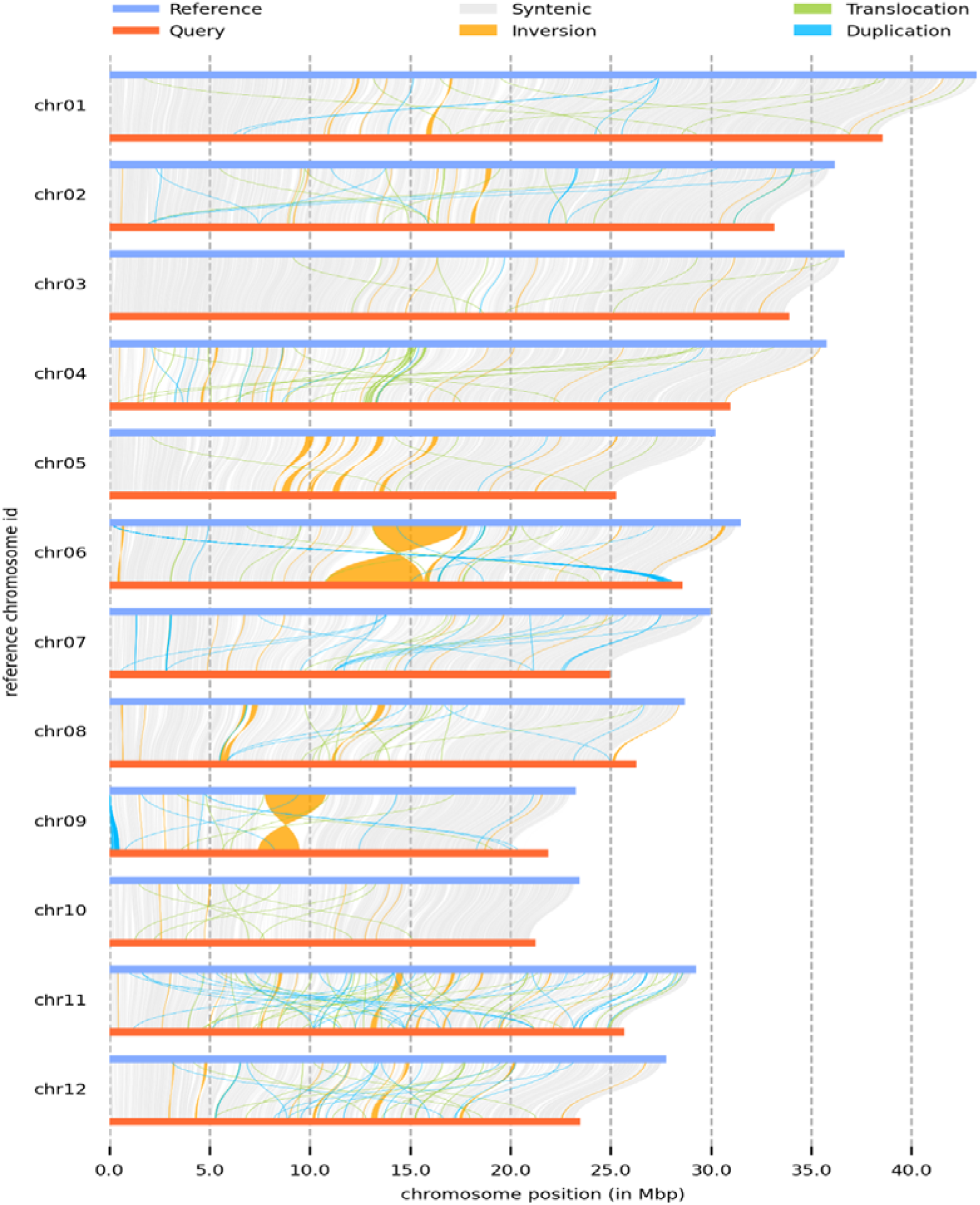
Collinearity and variation analysis of T2T genome of *O. longistatminata* and *O. sativa*. The reference genome being *O. satvia* and query genome being *O. longistatminata*.

### Analysis of SDs in the genome

Segmental Duplications (SDs) are genomic segments larger than 1 kb that repeat within the genome, exhibiting at least 90% sequence identity[16]. SDs frequently contain numerous duplicated genes, making them vital centers for gene’s innovation. Challenges in assembly technology made that the assembly of SD regions being collapsed, or entirely overlooked. As a result, this missing or inaccurate information limits our ability to understand the structure and evolution of a genome. The T2T genome of *O. longistaminata* offers an opportunity for more accurate characterization of SDs. In this study, we employed BISER to analyze the SDs in the rice genome[17], identifying 30.2 Mb of SDs, which constitute 9.12% of the genome. We discovered that SDs are not evenly distributed throughout the genome (Figure 3a). Instead, they are more frequently found on chromosomes 1 (chr1), 4 (chr4), 3 (chr3), and 2 (chr2), and less frequently on chromosomes 9 (chr9), 10 (chr10), and 5 (chr5). This distribution suggests that chromosomes 1, 4, 3, and 2 might have contributed to the evolution of rice in previously unrecognized ways (Table S8).

**Figure 3.**
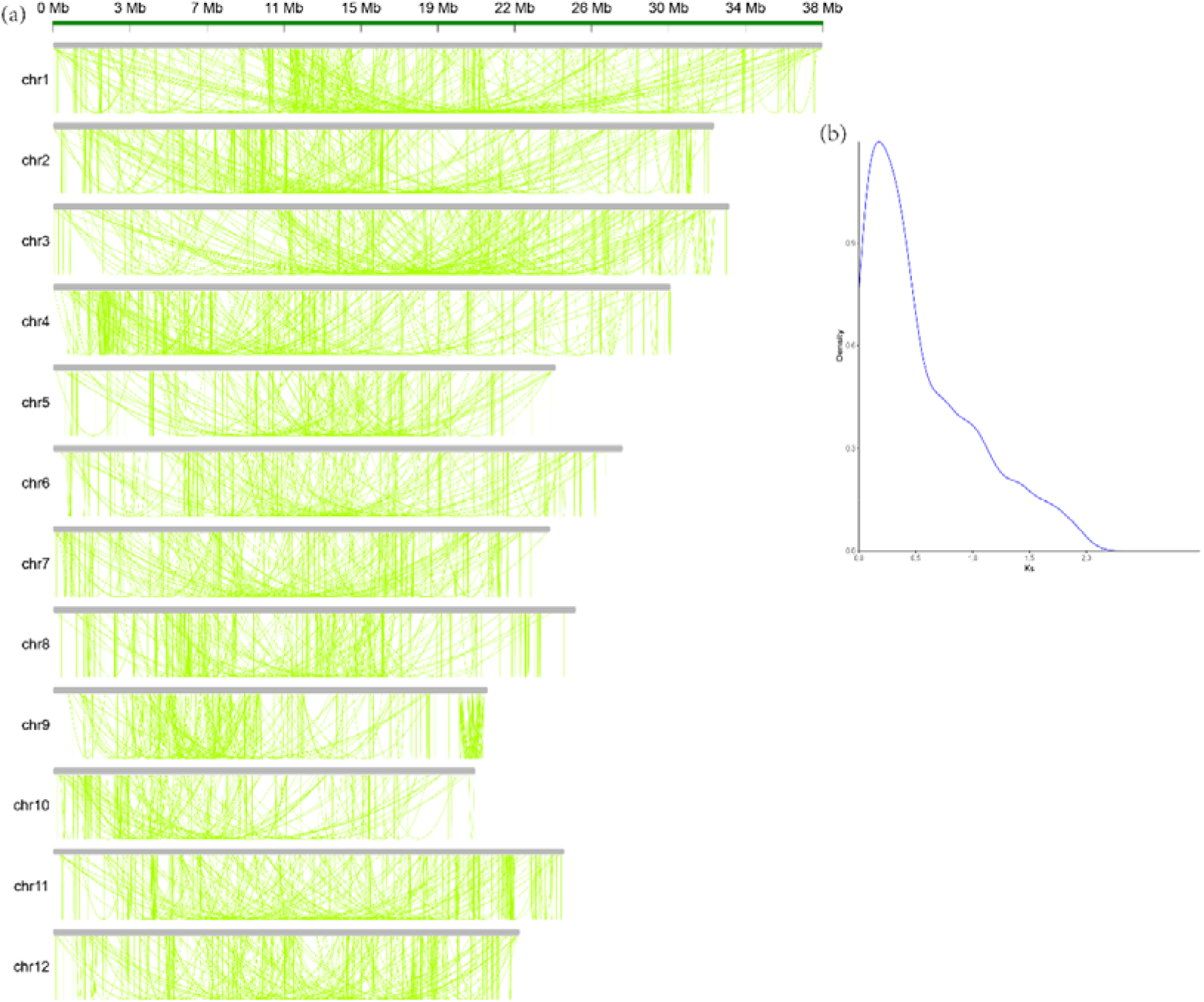
Segmental duplication analysis of the genome of *O. longistaminata*. (a). Distribution of intrachromosomal segmental duplication. (b) The density plot of the *Ks* value.

We proceeded to identify duplicated genes within the SD regions. Initially, we conducted an all-versus-all alignment using BLASTP[18] to identify potential paralogs, setting an E-value threshold of 10^−5^. In the SD regions, we identified a total of 4,179 pairs, of which 1,233 were the top matches for each other. For each paralogous gene pair within the SD regions, we calculated their *Ks* values as proxies for estimating the generation time of the corresponding SDs. Our findings indicate that the majority of these SDs were produced relatively recently (*Ks*=0.3) (Figure 3b). Gene ontology analysis showed that these genes were related to cellular amino acid metabolic processes, carboxylic acid metabolic processes and cofactor binding (Table S9).

### NBS gene family and transcription factors

Nucleotide-binding site-leucine-rich repeat (NBS-LRR) proteins as the largest family of resistance proteins are very important for plant’s defense against pathogens [19, 20]. We systematically investigated the NBS-LRR genes among 11 *Oryza* species (*O*.*barthii, O. brachyantha, O. glaberrima, O. glumipatula, O. indica, O. meridionalis, O. nivara, O. punctata, O. rufipogon, O. sativa, O. longistaminata*). There were 654 NBS-LRR genes in *O. longistaminata* genome, which were distributed in five different clusters (Table S10). Compared with other *Oryza* species, *O. longistaminata* has many fewer NBS-lRR domain genes, which reflects a contraction of resistance genes in this species. As NBS-LRR genes are essential components of the plant immune system, providing a mechanism for pathogen recognition and the activation of defense responses. The expansion of these types of genes in *O. longistaminata* may suggest that there was an increased ability for this species to adapt to its evolution.

We also investigated the variation in transcription factors among those *Oryza* genus species. For *O. longistaminata*, a total of 2095 transcription factors were distributed among 86 families (Table S11). The ERF transcription factor was the most abundant (857), followed by the bHLH family (128), NAC (120), MYB (119) and C2H2 (116). Intriguingly, we found that there were 47 FAR1 genes in *O. longistaminata* which was much larger than those in other African rice accessions. Research has shown that *FAR1* performs various functions across numerous cellular processes, indicating that *FAR1* is crucial for plant growth and development[21].

## Conclusion

In this study, we carried out a high-quality T2T assembly of wild rice *O. longistaminata*, which includes the complete assembly of 24 telomeres and 12 centromeres. We further compared this assembly with previously published *Oryza* genomes, and identified SVs between cultivated and wild rice accessions. Moreover, we investigated the SD genes, NBS-LRR resistance genes and transcription factors. This new *O. longistaminata* assembly represents a significant update, laying a fundamental evidential groundwork for focused investigations into genes associated with valuable phenotypic traits. It also sets the stage for future breeding endeavors, as well as further exploration into the evolutionary pathways of African rice and the *Oryza* genus.

## Methods

### Material preparation and sequencing

Fresh young leaves from mature *O. longistaminata* plants were collected from Yunnan University, Yunnan Province, China. Genomic DNA was extracted for Pacbio HIFI, MGI CycloneSeq and MGIseq. Pair-end libraries with 500-bp length were constructed and sequenced on the MGIseq platform. For Cyclone sequencing, genomic DNA was extracted by CTAB method[22], and the CycloneSEQ library was proceeded following the manufacturer’s guidelines. Each sample, comprising 2 μg of input DNA (≥21 ng/μL), was initially diluted with nuclease-free water to a total volume of 192 μL, followed by mixing with 14 μL DNA repair buffer 1, 14 μL of DNA repair buffer 2, 12 μL of DNA repair enzyme 1, and 8 μL of DNA repair enzyme 2. The mixtures were then incubated in a thermocycler through the following steps: 10 min at 20□, 10 min at 65□, and held at 4□. After incubation, the mixes were purified using a 1.0x volume of DNA clean beads, and DNAs were eluted with 240 μL of nuclease-free water. Next, the purified end-repaired samples were mixed with 10 μL sequencing adaptors, 100 μL 4x ligation buffer, 40 μL DNA ligase, and 10 μL of nuclease-free water before being incubated at 25□ for 30 minutes to complete the adaptor ligation. The ligated products were again purified with a volume of 1.0x DNA clean beads and long fragment wash buffer was applied to gently resuspend the beads. After removing the supernatant, the libraries were recovered in 42 μL of elution buffer and quantified on a Qubit fluorometer. Each prepared library was sequenced on the CycloneSEQ WuTong02 platform according to the protocol. A total of 25.6 Gb of clean subreads with a length longer than 79 Kb were obtained and used as Ultra-long reads. For the construction of the PacBio HiFi library, more than 5 µg of DNA was prepared for size selection using a BluePippin instrument. Subsequently, PacBio Sequel II single-molecule real-time (SMRT) bell libraries of approximately 20 kb were constructed in accordance with the PacBio protocol. The library was loaded into SMRT cells with the DNA Sequencing Reagent Kit. These SMRT cells were then run on a PacBio Sequel II CCS system, which generated 24 Gb of long-read sequencing data.

### Genome assembly

We employed hifiasm (version 0.19.5-r592) for genome assembly, utilizing both HIFI reads and ultra-long CycloneSeq reads under the mixed assembly model with default settings[23]. Subsequent polishing of the assembled genome was done using NextPolish[24] with MGISEQ reads. For chromosomal anchoring of the contigs, we first utilized cleaned HiC reads. Unique mapping reads were identified using bowtie2 (v 2.3.2) [25], followed by the detection of valid interacting paired reads via Juicer (v 2.8.1) [26]. These valid read pairs were then used to construct pseudo-chromosome sequences with 3D-DNA[27]. The HiC interactions are shown as heat maps through Juicebox. After that, the genome had only one gap. Finally, gap filling was accomplished by using TGS-Gapcloser (v 1.2.1)[8] with CycloneSeq reads, and corrections were made by using pilon (v 1.24) with MGI paired-end reads[28]. Based on the genome of *O. sativa*, we artificially reoriented some chromosomes and renamed their chromosome numbers.

To access the quality of the assemblies, we mapped the short paired-end reads to the assembly by using the bwa-mem tool from BWA [29], and BUSCO analysis was performed with the embryophyte_odb9 database[11].

### Genome annotation

We initially generated repetitive libraries for both species through a dual approach involving homology comparison and de novo prediction. Repetitive sequences were identified using LTR Finder[30] and RepeatModeler [31]. Homology-based prediction was performed using TRF[32] and RepeatMasker[33] with the Repbase TE library. The annotated and classified repetitive sequences were then used to mask the genomes with RepeatMasker. Additionally, we employed LTR Retriever[34], in conjunction with LTR Finder, to calculate the LTR Assembly Index (LAI), which assesses assembly continuity by evaluating the assembly of repeat sequences[12].

For the RNA-Seq assisted predictions, ISO-seq sequences were obtained from a mixed tissue. We employed SMRT Link v8.0, applying the parameters –min-passes 3, –min-length 50, –max-length 15000, and –min-rq 0.99, to refine the circular consensus sequence (CCS) subreads, subsequently gathering high-quality reads. For classifying the full-length reads, Lima v2.2.0 was utilized with the following settings: –isoseq, –dump-clips, and –peak-guess. The final assembly of full-length Iso-seq transcripts was achieved using isoseq3[35], which employs the refine module (parameters: –require-polya and –min-polya-length 20) and the cluster module (parameters: –verbose and –use-qvs). After that, Transdecode[36] was used to predict the CDS and the longest CDS sequences were fed into Maker for gene annotation. For homologous predictions, protein sequences from *Zea Mays, O. sativa*, and *A*.*thaliana* were used. MAKER2 pipeline [37] was used for protein-coding genes annotation, and de novo gene models were accessed by AUGUSTUS[38] and Fgenesh [39]. To predict gene functions, we performed a BLAST search of their protein sequences against the Swiss-Prot and NR databases, using a threshold E-value of 1e-5. Subsequently, we employed InterProScan to annotate motifs and domains by searching for matches in those databases.

### Structural variation analysis

We utilized a suite of tools from MUMmer4[40] to analyze genomic differences between *O. longistaminata* and *O. sativa*. The nucmer tool was employed to compare syntenic chromosomes, and the results were subsequently filtered through a delta-filter with the parameters ‘-c 100 -b 500 -l 50’. The alignments were then converted into tab-delimited files using the show-coords program. Finally, SyRI was applied to identify structural variations (SVs) [41].

### Identification of telomeres and centromeres

In most plants, telomere sequences consist of short, conserved satellite repeats arranged in tandem. We identified a typical plant telomere sequence (CCCTAAA at the 5’ end or TTTAGGG at the 3’ end) and subsequently identify telomeres across all 12 chromosomes. To detect centromeric regions, we employed the quarTeT tool[10]. This tool is well-suited for identifying genomic areas with high and low gene densities, as well as short tandem repeats, which are characteristic features of centromeric regions.

### Detection of Segmental Duplications (SDs)

Briefly, our genome assembly underwent an initial soft-masking process, during which all common and tandem repeats were converted into lowercase letters. Following this, BISER was employed to detect Segmental Duplications (SDs), utilizing its default parameters[17].

### *Ks* of the duplicated gene pairs

Protein sequences from duplicated gene pairs within SDs were extracted and subsequently aligned utilizing the MUSCLE alignment program[42]. These aligned protein sequences were then transformed into corresponding coding sequence alignments through PAL2NAL[43]. Next, we calculated the rate of synonymous substitutions per synonymous site (*Ks*) for each gene pair, employing the KaKs_Calculator[44]. The distribution of *Ks* values was graphically plotted and visualized using the R statistical software.

### Identification of *NBS-LRR* genes and TF genes

For each species analyzed, protein sequences were extracted and subsequently screened against the raw Hidden Markov Model (HMM) of the NB-ARC family (PF00931) utilizing HMMER (version 3.1b1), applying the default parameters[45]. NBS-specific HMMs were constructed utilizing the hmmbuild program within HMMER, which were then employed to identify NBS-encoding proteins. To identify specific protein domains, PfamScan was utilized for screening these proteins against the Pfam-A database (Pfam31.0) [46]. Additionally, coiled-coil domains were detected using the ncoils tool [47], with its default parameters applied.

## Acknowledgements

This work was supported by the Key Laboratory of Genomics, Ministry of Agriculture, Guangdong Provincial Key Laboratory of core collection of crop genetic resources research and application, Shenzhen Engineering laboratory of Crop Molecular design breeding, the National Natural Science Foundation of China (32322063 to Shilai Zhang) and the Shenzhen Science and Technology Program (KQTD20221101093603011 to Jingnan Yang). This work is a part of the 10KP project, and is also supported by the China National GeneBank.

## Authors’ contributions

WT. and Y.D. conceived the project and were responsible for the project initiation. J.Y. and Y.D. designed the experiments. S.Z., H.Liu, X.L., and F.H. contributed to sample preparation. F.G., T.Z., H.Lu, C.C., F.L.,Z.L.,Y.H., W.L., L.Y.,X.X., and Y.G. performed the experiments and sequencing. X.G., L.L., D.F., J.Y. and W.H. performed the data analysis. X.G. wrote the original draft. T.W., S.K.S., and Y.D. revised the manuscript. All authors read and approved the manuscript.

## Data availability

The T2T genome assembly and sequenced Cyclone reads has been deposited to the CNSA (CNGB Nucleotide Sequence Archive) with accession CNP0005176 (https://db.cngb.org/cnsa/).

