## Supplementary Figures for "Telomere-to-telomere African wild rice (*Oryza longistaminata*) reference genome reveals segmental and structural variation"

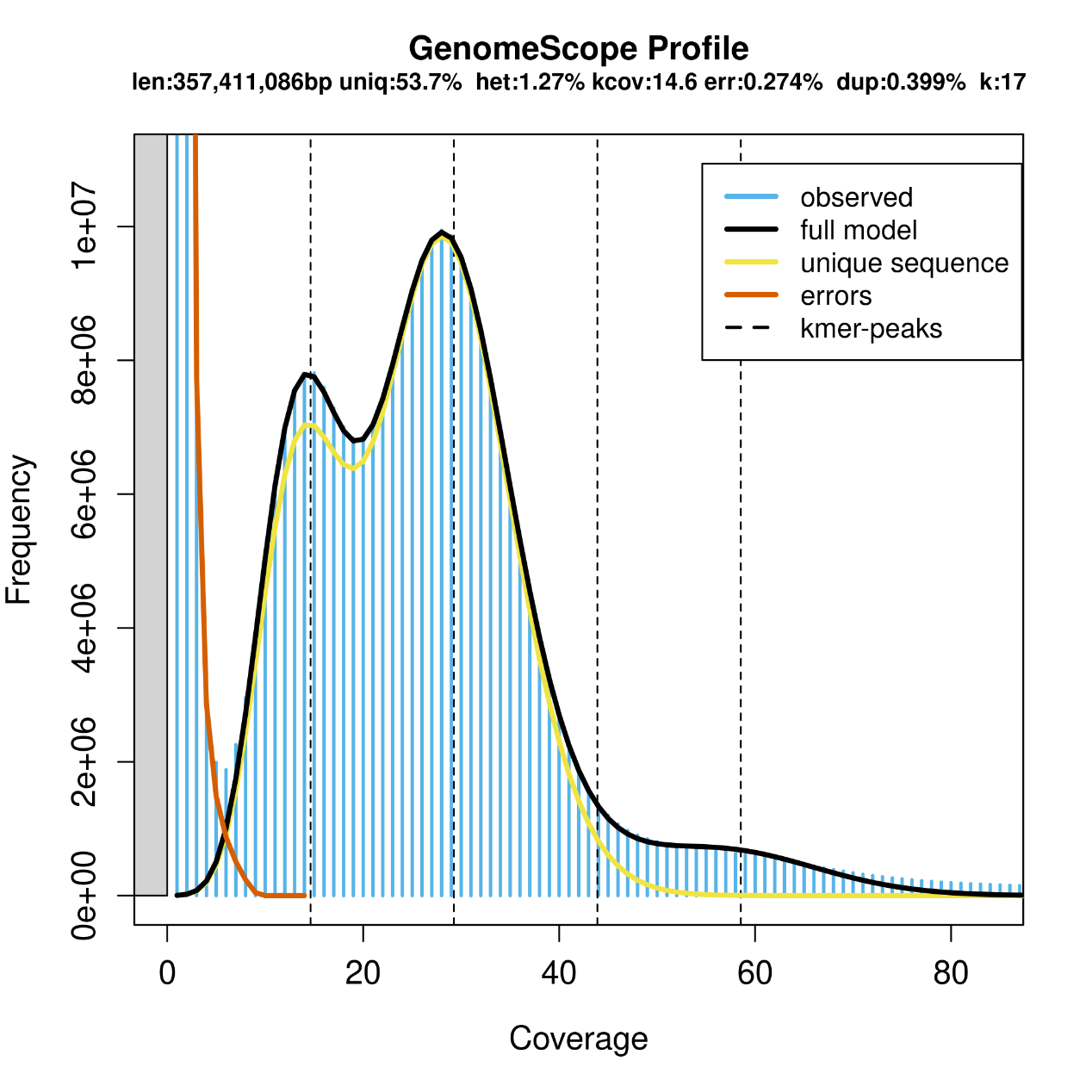

**Figure S1. *K*-mer distribution of the *O.longistaminata* genome. *K*-mer=17.**

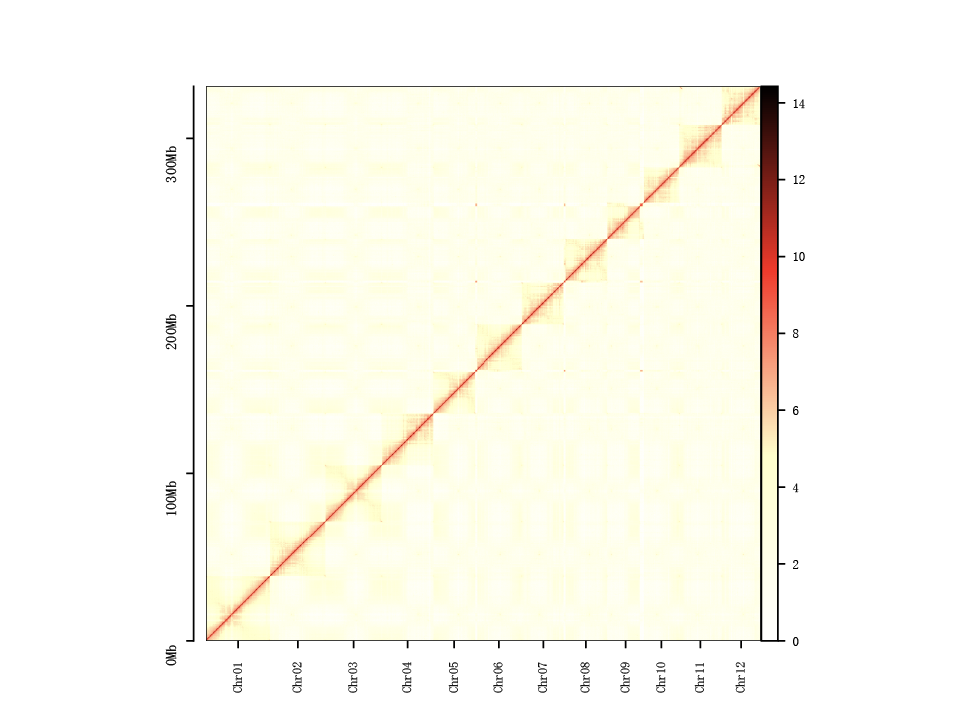

**Figure S2. Heatmap for Hi-C assembly of *O.longistaminata* genome.**

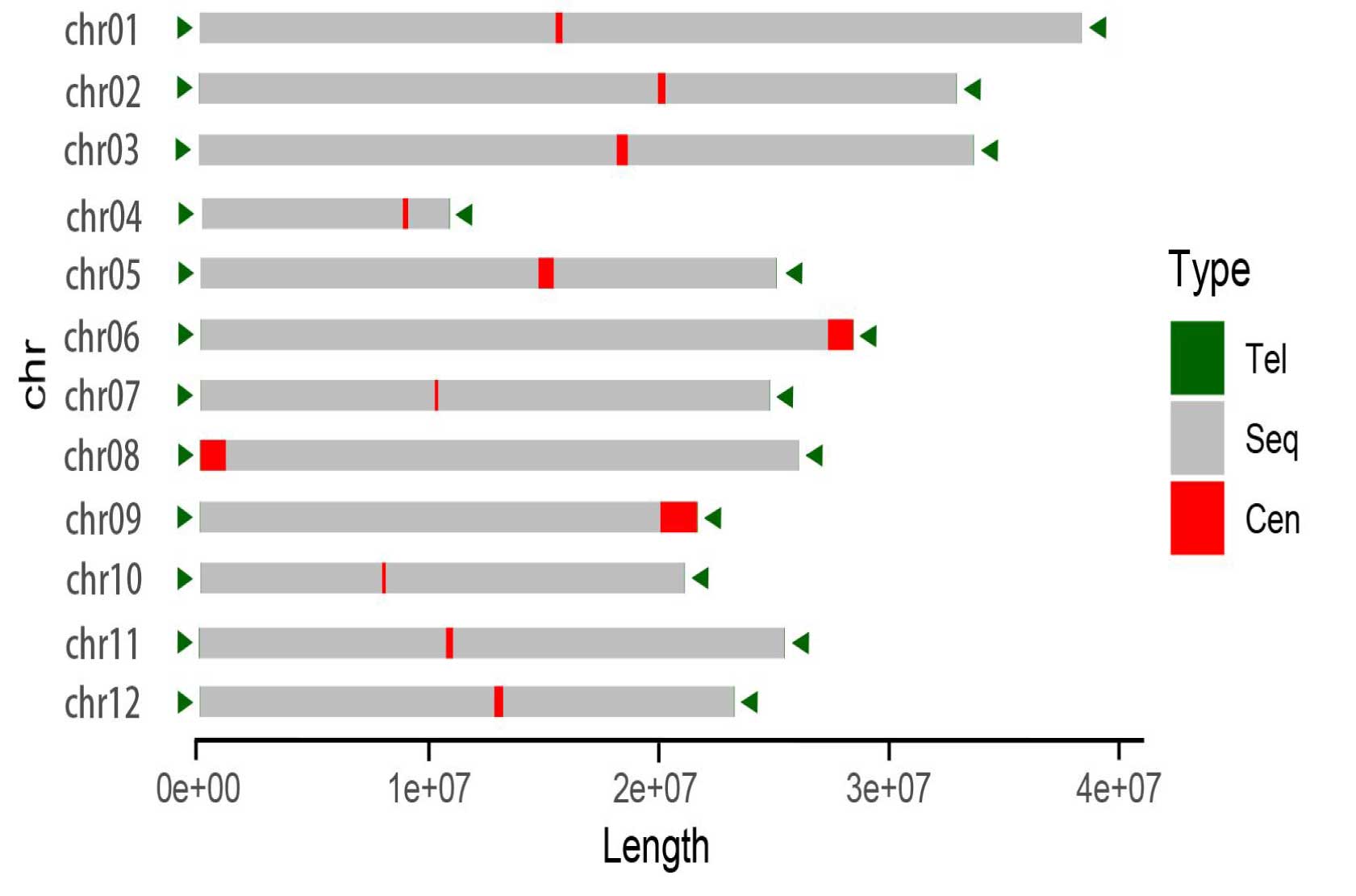

**Figure S3. The overview of the genome structure.**

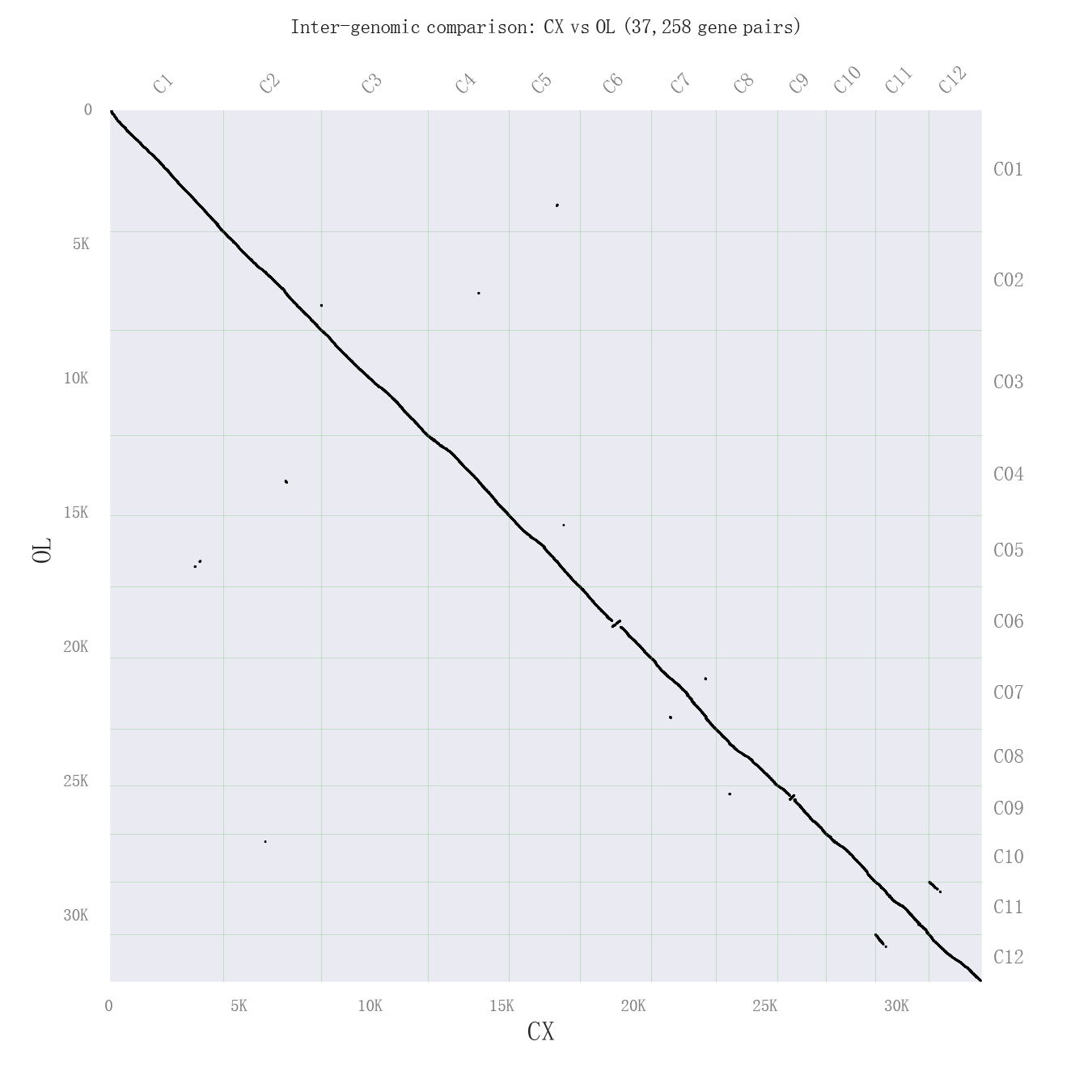

**Figure S4. Homologous dot-plot between T2T genome and previous *O.longistaminata* genome genomes.**

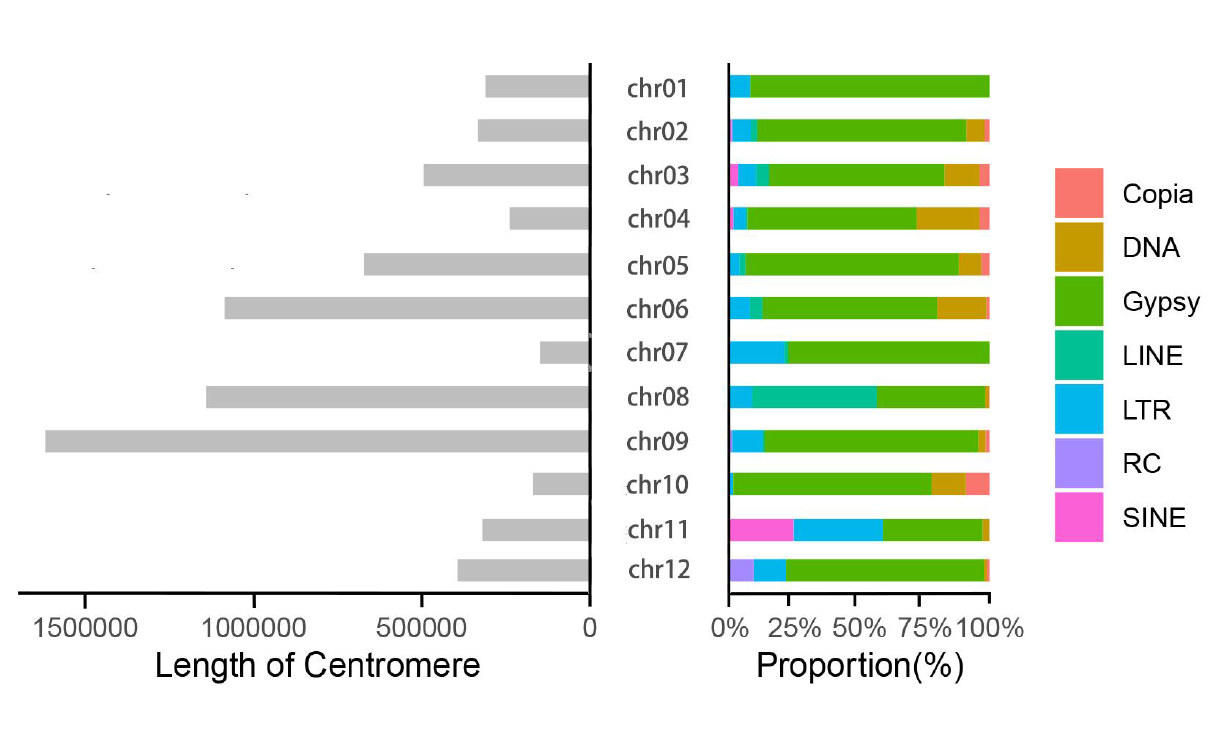

**Figure	S5. The length of Centromere and the repeat content of its distribution**
